# Adult fibroblasts retain organ-specific transcriptomic identity

**DOI:** 10.1101/2021.06.03.446915

**Authors:** Elvira Forte, Mirana Ramialison, Hieu T. Nim, Madison Mara, Rachel Cohn, Sandra L. Daigle, Sarah Boyd, J. Travis Hinson, Mauro W. Costa, Nadia A. Rosenthal, Milena B. Furtado

**Affiliations:** The Jackson Laboratory, Bar Harbor, ME 04609, USA; The Australian Regenerative Medicine Institute, Monash University, Clayton, VIC 3800, Australia; Systems Biology Institute Australia, Clayton, VIC 3800, Australia; Murdoch Children’s Research Institute, Parkville, VIC 3052, Australia; The Jackson Laboratory, Farmington, CT 06032, USA; Centre for Inflammatory Diseases, School of Clinical Sciences at Monash Health, Monash University, Clayton, VIC 3168, Australia; National Heart and Lung Institute, Imperial College London, London SW72BX, UK

## Abstract

Organ fibroblasts are essential components of homeostatic and diseased tissues. They participate in sculpting the extracellular matrix, sensing the microenvironment and communicating with other resident cells. Recent studies have revealed transcriptomic heterogeneity among fibroblasts within and between organs. To dissect the basis of inter-organ heterogeneity, we compare the gene expression of fibroblasts from different tissues (tail, skin, lung, liver, heart, kidney, gonads) and show that they display distinct positional and organ-specific transcriptome signatures that reflect their embryonic origins. We demonstrate that fibroblasts’ expression of genes typically attributed to the surrounding parenchyma is established in embryonic development and largely maintained in culture, bioengineered tissues, and ectopic transplants. Targeted knockdown of key organ-specific transcription factors affects fibroblasts functions, with modulation of genes related to fibrosis and inflammation. Our data open novel opportunities for the treatment of fibrotic diseases in a more precise, organ-specific manner.

## Introduction

Fibroproliferative disorders are the main cause of mortality and morbidity in the developed countries, accounting for about 45% of deaths in the United States [1]. Despite the impactful prevalence of chronic organ fibrosis, current anti-fibrotic drugs are both inefficient and non-specific to this condition [2, 3]. Fibroblasts, main players in fibrosis, have gained increased attention for their capacity to provide functions far beyond their canonical secretion of extracellular biological scaffolding and formation of scar tissue after injury. Recent literature poses the organ fibroblast as a major regulatory hub that senses local microenvironment imbalances and controls tissue remodeling [4] upon activation and phenotypic differentiation into the pro-fibrotic myofibroblast [5]. They are also involved in immunomodulation [6], by producing and responding to cytokines and activate immune cells of the innate and adaptive immune systems [7, 8], through organ-specific regulatory networks [9].

Organ fibroblasts have been historically difficult to identify and study *in vivo*, due to their vague functional definition and lack of adequate markers that label organ fibroblast pools completely and specifically [10]. Recent advances in lineage tracing and multiomics single-cell analyses have revealed a significant heterogeneity of fibroblasts within and among tissues, and we are just beginning to understand how fibroblast heterogeneity corresponds to distinct functions [3, 11–14]. Despite being morphologically similar, spindle-shaped mesenchymal cells located in stromal tissues, fibroblasts acquire specialized functions related to their anatomical position [9, 15, 16] and appear to retain a positional memory of the embryonic developmental axis: anterior-posterior, proximal-distal and dermal non-dermal, possibly reflecting their role in conveying positional identity in embryogenesis [17–20], suggesting responsiveness to molecular cues that drive body compartmentalization. Fibroblast heterogeneity within an organ tends to arise from the distinct embryological origin and/ or anatomical localization [12, 14, 21, 22], while inter-organ differences have been mostly ascribed to the matrisome, as shown by the transcriptomic comparison among fibroblasts from muscular tissues [21].

Having previously reported that fibroblasts isolated from the adult mouse heart retain a cardiogenic transcriptional program [23], we show here that fibroblasts isolated from different adult organs similarly retain the expression of transcription factors and other gene sets involved in the determination of organ formation and patterning during embryonic development. This signature is captured in nascent embryonic organ fibroblasts, retained in isolated adult cultured cells, in co-culture with parenchymal cells from different organs, or when ectopically transplanted into a different organ *in vivo*, despite adaptation to the new environment. The robustness of the fibroblast organ transcriptome signature under different microenvironmental challenges supports its importance for organ interaction, connectivity and function. In addition, knock-down of selected organ development transcription factors in cardiac fibroblasts de-regulated the expression of genes involved in inflammation, fibrosis and ECM deposition, further supporting the relevance of these genes in fibroblasts function. In summary, our study uncovers stable expression of organ-specific, development-related signature genes in adult fibroblasts, thus offering new prospects for possible targeted anti-fibrotic therapies.

## Results

### Metabolic and extracellular matrix components comprise a generic fibroblast gene signature

To compare the gene signature of fibroblasts from different organs and eliminate potential RNA contaminants from other organ cell types, dissociated adult murine tissues were cultured for five days, followed by sorting for CD45-CD31-CD90+ fibroblasts [23] (**Supplementary Fig. 1)**. High-throughput gene expression profiling identified 1281 highly expressed genes common to all fibroblast types, comprising the generic fibroblast signature (**Supplementary Table 1**).

Through Ingenuity Pathway Analysis (IPA, Qiagen), we classified the commonly expressed genes based on cellular function (**Supplementary Fig. 2a)**, and cellular localization (**Supplementary Fig. 2b**). Top functions included mechanisms of cell maintenance, such as proliferation, cytoskeletal arrangement and cell movement, as well as general metabolic processes, including carbohydrate, nucleic acid protein and small molecule biochemistry. Common fibroblast identifier genes were encountered within various IPA process classification groups. As an example, the cell surface receptor CD90 (*Thy1* gene) belongs with cellular assembly and organization, growth and proliferation and protein synthesis, while the myofibroblast marker smooth muscle actin (*Acta2* gene) was found in functions of cellular movement. The presence of CD90 and absence of CD31 (*Pecam1* gene)/CD45(*Ptprc* gene) in all organ groups validated our positive/negative selection strategy for cell isolation, indicating a generally consistent population of cells in all organs. Extracellular matrix (ECM) elements, including collagens, were included in several functional annotations, such as to cell morphology, assembly and organization, cellular compromise, function and maintenance or cell signaling.

In terms of cellular localization, most fibroblast identifier genes encoded proteins localized to the cytoplasm (**Supplementary Fig. 2b, Supplementary Table 1**). Fibroblasts are generally defined as ECM secreting cells. Common ECM genes consistently present amongst fibroblast samples included collagens *Col1a1*, *Col1a2*, *Col3a1*, *Col4a3* and *Col5a1* (**Supplementary Fig. 2b**). Other extracellular genes encoded members of secreted *Tgfb*, *Fgf, Ins, Igf1 Vegf, Egf*, *Notch* and *Wnt* pathways (**Supplementary Table 1**). Among membrane genes were integrins *Itga3/9*, *Itgb3/6*, collagen receptor *Ddr1*, and importantly, receptors for adrenalin (*Adrb1*), acetylcholine (*Chrm1*), angiotensin II (*Agtr1, Agtr2*) and calcium (*Cacna1c*) and potassium (*Kcnab1* and *Kcnj11*) channels. The nuclear and cytoplasmic gene fractions included mostly housekeeping genes, but also *Acta2* and fibroblast markers filaminA (*Flna*) and vimentin (*Vim*). The small fraction of genes not classified by the IPA cell compartment analysis, including the growth factor neuregulin 1 (*Nrg1*), were placed in the category “Other”.

### Organ fibroblasts retain *HOX* codes

The *HOX* code defines body segmental identity and is highly conserved from flies to mammals. *HOX* genes show colinear expression and undergo chronological activation in the embryo, where upstream genes successively activate downstream genes in an antero-posterior fashion, such that upstream genes are activated first in more anterior segments of the body. In mammals, the *HOX* cluster has undergone a series of duplications and deletions that led to the formation of four paralogous clusters a, b, c and d (**Fig. 1a**).

**Fig. 1:**
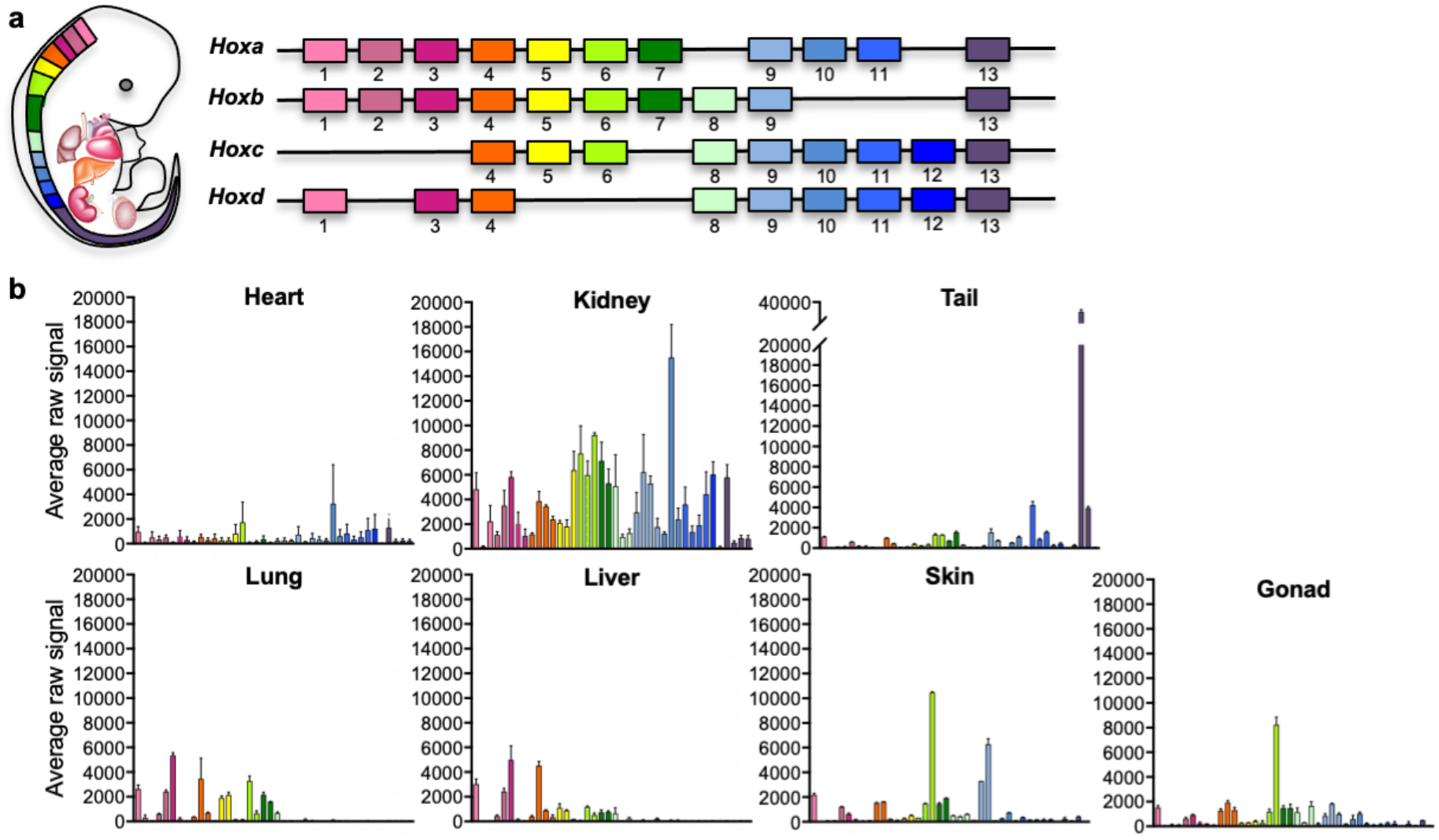
Positional Code of Organ Fibroblasts. **a** Murine Hox cluster code, showing proximally expressed Hox genes in pink and distal ones in purple. **b** Bar plots showing the average raw signal for all expressed *Hox* genes in organ-specific fibroblast samples. Data are mean ± SEM of 3 biological replicates for each fibroblast type.

Site-specific *HOX* expression has been previously reported in human skin fibroblasts [17, 18, 20] and mouse mesenchymal cells isolated from different organs [19], and it has been shown to be cell-autonomous and epigenetically maintained, suggesting a source of positional memory to differentially pattern tissue-specific homeostasis and regeneration. To determine if fibroblasts isolated from other adult organs retain a distinct *HOX* signature, we plotted the average raw expression of all *HOX* genes per each fibroblast type (**Fig. 1b** **and Supplementary Figure 3, Table 2**). Among profiled organs, five patterns of *HOX* expression were identified: lung and liver showed expression of anterior *HOX* genes, in particular genes from the clusters 1 to 7, although liver had lower *HOX 4-7* expression when compared with lung. A second group including skin and gonad displayed high *HOX 6* expression, with skin from thoracic and abdominal ventral skin areas also expressing high *HOX 9* gene levels. The third classification group was represented by the heart, characterized by low *HOX* gene expression. This may reflect embryonic developmental processes, as *HOX* genes are known to exert minimal influence on heart formation, and are generally not expressed in the heart, except for the residual expression carried over by neural crest cells that invade the arterial pole of the heart and promote aorticopulmonary septation [24]. The great vessels were excluded from our sample collection, and therefore cells of neural crest origin were likely not captured in the analyses. The fourth classification group was represented by kidney, expressing intermediate to high levels of most anterior *HOX* genes up to *HOX11*, consistent with previous observations for the developing kidney [25]. The fifth category, represented by the mouse tail, had a posterior *HOX* code signature, represented by *Hox13*, which correlates with previous findings for human distal segment fibroblasts, represented by feet skin fibroblasts [17, 18]. Taken together with previous observations, these analyses confirm that adult organ fibroblasts retain positional *HOX* gene expression signatures, generally reflecting the embryological segmental identity of organ fibroblasts.

### Organ fibroblasts show unique molecular signatures

To highlight the unique transcriptomic signatures of these positionally distinct fibroblast pools, we performed a differential expression analysis and considered genes that were enriched by 10-fold change or more in single organ fibroblasts compared to tail fibroblasts (**Fig. 2, Supplementary Table 3**). Gene Ontology annotation revealed organ development programs; processes such as epithelial development, hepatoblast differentiation, lung lobe development, kidney development, reproductive process and heart development were found enriched in their respective organ fibroblast pools (**Fig. 2a**). Strikingly, signature embryonic transcription factors, i.e., genes with established involvement in organ development, were enriched in organ-specific subsets, including *Tbx20*, a crucial transcription factor for heart development previously described in cardiac fibroblasts [23]. Likewise, genes essential for lung morphogenesis (*Foxf1)*[26], liver development (*Hhex*) [27], early kidney formation (*Pax8)* [28] and gonad development *(Lhx9)* [29] were all specifically enriched in fibroblasts from their respective organs. Expression of signature genes was validated by qPCR (**Fig. 2b, Supplementary Fig. 4**) and immunocytochemistry (**Fig. 3**). In general, signature gene expression patterns in embryonic fibroblasts were retained in fibroblasts from adult tissues (*Krt4*, *Krt6a*, *Serpinb5* and *Hp* for skin; *Tbx20* and *Col2a1* for heart; *Foxf1* for lung; *Hhex* and *Foxa2* for liver, *Bmp7* and *Pax8* for kidney, *Cyp11a1* and *Lbx9* for gonad). Significant expression of *Hhex* and *Bmp7* were also found in several organs during embryonic development but were restricted to a single organ in adulthood. As an exception to single organ enrichment, *Foxa2* was also substantially upregulated in lung fibroblasts (∼20 fold), in addition to liver fibroblasts (∼30 fold).

**Fig. 2:**
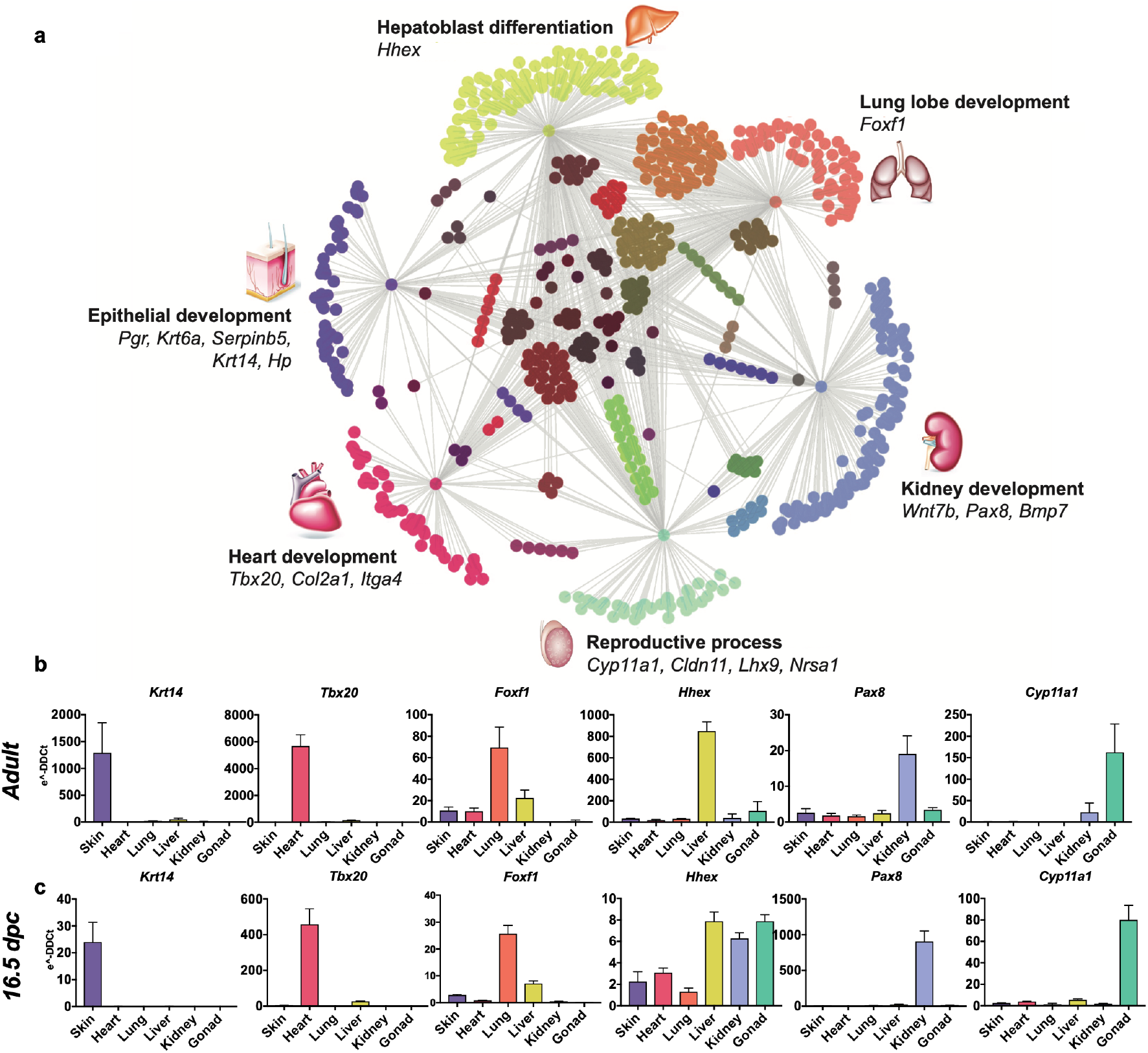
Embryological Molecular Signature of Organ Fibroblasts. **a** Cytoscape representation of network of genes (dots) singularly expressed in organ fibroblasts (only one grey edge between the gene and the organ) or shared among organs (multiple grey edges linking the gene to several organs). Genes involved in organ development are highlighted. **b-g** Validation of the expression of selected organ-enriched, developmental related genes using qPCR on cultured organ-derived fibroblasts isolated from adult mice (top row) or E16.5 embryos (bottom row). Data are mean ± SEM of the e^-DDCt values, on 3 biological replicates. The housekeeping gene is *Hprt* and the reference sample is tail fibroblasts.

**Fig. 3:**
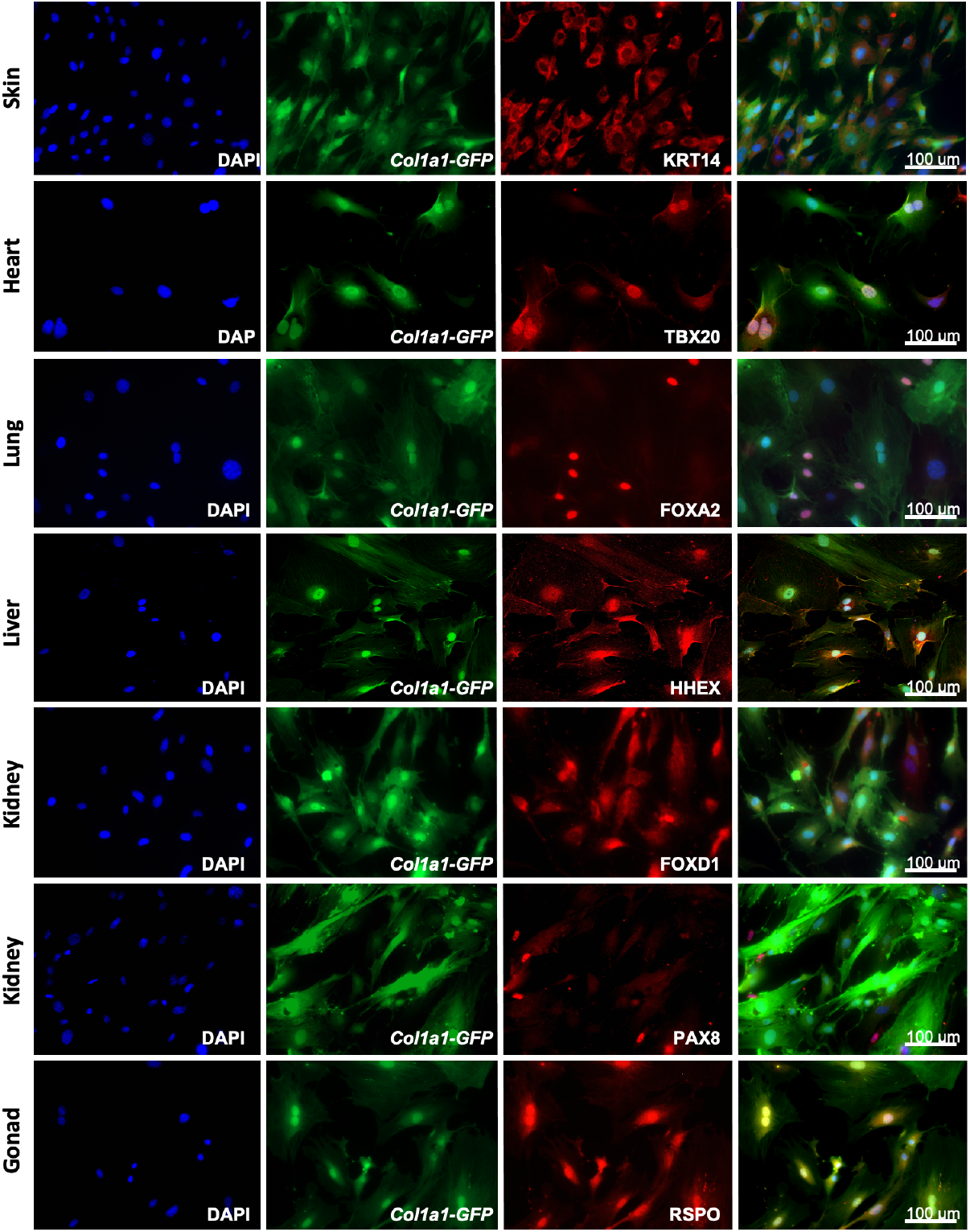
Adult fibroblasts express organ specific transcription factors. Immunocytochemistry for organ specific markers (KRT14, TBX20, FOXA2, HHEX, FOXD1, PAX8, RSPO) on adult fibroblasts obtained from different organs of *Col1a1-GFP* mice and cultured for 5 days. Scale bar =100um.

IPA analysis delineated top canonical pathways, diseases, functions and networks associated with selectively enriched genes in each fibroblast populations and supported the argument that fibroblasts retain molecular identity of their organ developmental origins (**Supplementary Figs. 5-10**). Among organ-related processes enriched in fibroblast subsets were dermatological diseases and conditions and morphogenesis of the epithelial tissue for skin fibroblasts (**Supplementary Fig. 5**), respiratory system development for lung fibroblasts (**Supplementary Fig. 6**), liver development for liver fibroblasts (**Supplementary Fig. 7)**, acute renal failure, metanephros development and kidney formation, and abnormal kidney development, disease and function for kidney fibroblasts (**Supplementary Fig. 8**), reproductive system development, function and disease, morphology of genital organs and primary sex determination networks and reproductive system dysfunction for gonad fibroblasts (**Supplementary Fig. 9**), cardiovascular disease development and function, cardiac enlargement and disease and cardiac developmental processes for heart fibroblasts (**Supplementary Fig. 10**).

To establish the translational relevance of our findings, the presence of 42 genes uniquely enriched in cardiac fibroblasts (log2, 10-fold, FDR <0.01) was determined in left ventricular heart biopsies from healthy (N=5) and chronic ischemic heart failure patients (N=5) (**Supplementary Fig. 10g, Table 4**). Ischemic heart failure was chosen for analysis due to the likelihood of replacement fibrosis as a pathological signature. Out of the 42 murine cardiac fibroblast genes, 28 were present in both control and heart failure samples, including *Tbx20*, which was unchanged between control and heart failure. *FNDC1*, *FRZB*, *MFAP4* and *OLFML1* were significantly up-regulated in ischemic heart failure; while *MFSD2A*, *PNP* and *SERPINA3N* were down-regulated. Four genes had no human homolog and ten were not found in the human heart transcriptome dataset. These findings confirm commonalities across species and further identify potential candidates of fibrotic interest for future investigation.

### Organ-enriched gene expression is retained at the single cell level in freshly isolated and cultured fibroblasts

Single-cell RNA seq (scRNAseq) is a powerful tool to determine granularity of gene expression at the population level. To assess how organ signatures are reflected in freshly isolated fibroblasts, we re-analyzed the stromal cell dataset from a publicly available multi-organ single-cell RNA seq (scRNAseq) study (the Mouse Cell Atlas) [30]. Focusing on lung, testis, kidney, liver and neonatal heart cells, we unbiasedly identified 8 populations, including 3 lung and 2 kidney sub-clusters (**Fig. 4a**). Pairwise differential expression analysis supported a previously reported classification of lung fibroblasts populations [31], with two types of matrix fibroblasts - “LungA” (Col*14a1, Pi16, Dcn* enriched), “LungB” (*Col13a1, Cxcl14, Tcf21* enriched)- and a group of myofibroblasts- “LungC” (*Acta2*, *Myl9*, *Tagln* positive) (**Supplementary Fig. 11 a-b)**. For kidney clusters, “KidneyB” showed higher levels of canonical fibroblast markers *Dcn*, *Gsn* and *Col1a2*, while the larger “KidneyA” expressed relatively higher levels of genes involved in response to injury, or in renal carcinoma metastasis and progression (*Spp1, Krt8*), suggesting that this cluster is composed of tubular cells acquiring a mesenchymal phenotype *in vivo* [32] (**Supplementary Fig. 11 c-d)**, as kidney epithelial cells are known to undergo dedifferentiation *in vivo* and *in vitro* to repair tubular injuries [33, 34].

**Fig. 4:**
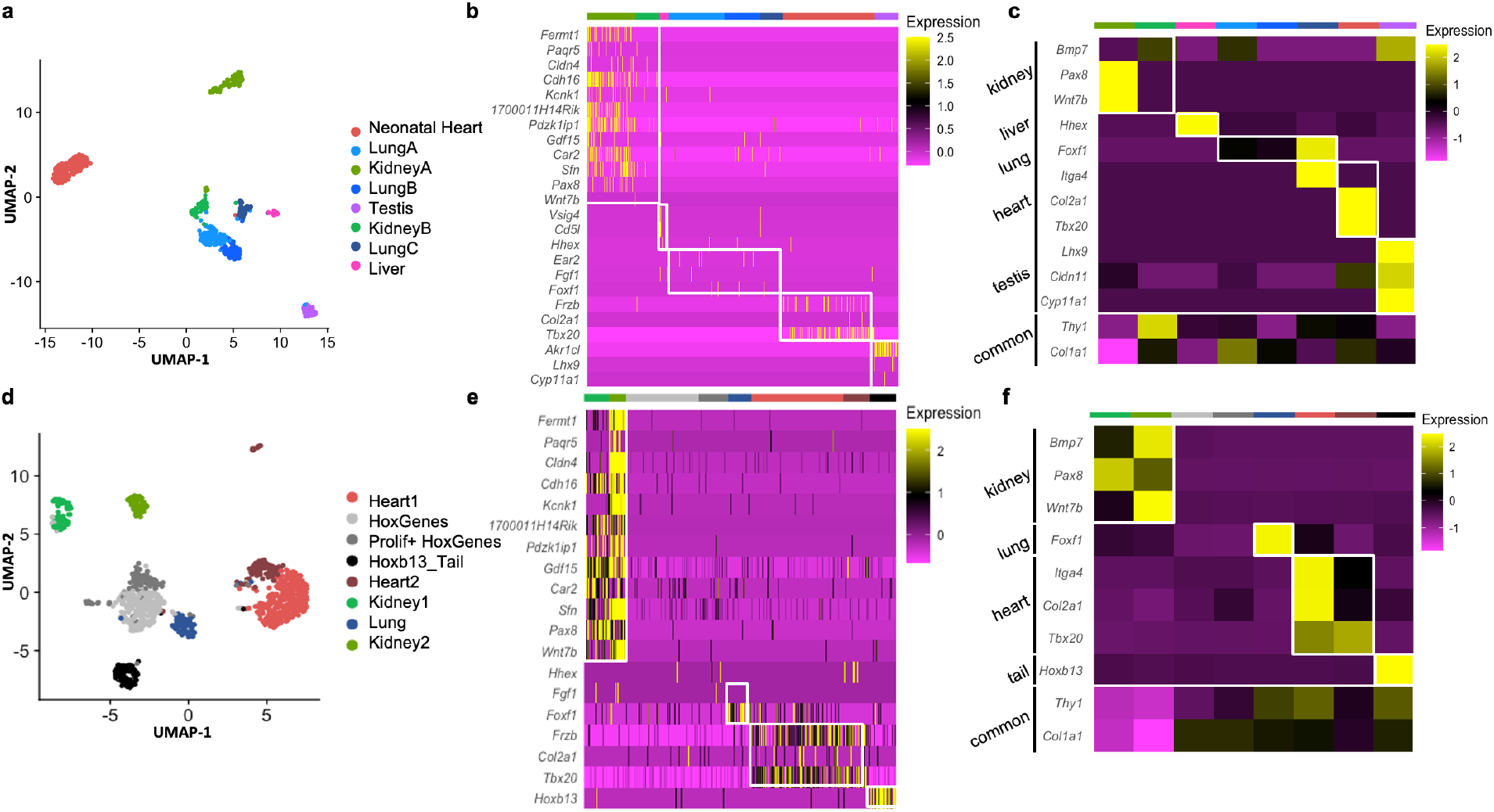
Analysis of organ specific signatures at the single cell level. **a-c** Re-analysis of the Mouse Cell Atlas stromal cell dataset. **a** UMAP visualization of selected stromal cell populations (682 cells). **b** Heatmap representing the expression of top-organ specific genes, identified from the bulkRNAseq comparison, on individual cells in the scRNAseq dataset. **c** Heatmap showing the average expression per population of organ-development related genes (same as shown in Fig.2). **d-f** scRNAseq analysis of mixed cultured stromal cells of different origins. **d** UMAP visualization of the captured cells. **e** Heatmap representing the expression of top-organ specific genes, identified from the bulkRNAseq comparison, on individual cells. **c** Heatmap showing the average expression per population of organ-development related genes (same as shown in Fig.2).

Overall, the expression of organ-specific (**Fig. 4b**) and development-related genes (**Fig. 4c**) previously identified in the bulk cultured cell analysis was preserved, despite the reduced coverage and expected heterogeneity at the single cell level (**Fig. 4b**). Interestingly, when multiple subclusters were present (as in lung and kidney fibroblasts) the expression of the organ-specific genes was enriched in myofibroblasts (Lung C expressed relatively higher levels of *Foxf1*) or activated fibroblasts (Kidney A, expressed higher levels of *Pax8* and *Wnt7b)* (**Fig. 4c, Table 5, Supplementary Fig. 11**).

Further confirmation of the organ enriched program was obtained with scRNAseq of pooled primary cultures from different origins (kidney, liver, lung, heart, skin, testis, tail) (**Fig. 4d**). Unbiased clustering defined Kidney 1-2, Lung, Heart 1-2, and Tail fibroblasts. Two additional clusters were unclassified based on organ identity, although marked by the expression of Hoxc genes (*HoxGenes*) and proliferation genes (*Prolif+HoxGenes*) (**Fig. 4d-f, Table 6**). Highly expressed (**Fig. 4e**) and development-related genes (**Fig. 4f**) from original bulk analysis were again confirmed in these organ populations. Both cultured kidney clusters (Kidney 1-2) expressed the epithelial stress response marker (*Spp1*) and were transcriptionally closer to freshly isolated “KidneyA” (**Supplementary Fig. 12c**), possibly representing two stages of tubular cells epithelial-to-mesenchymal transition [35]: Kidney1 had higher expression of myofibroblast genes (*Col4a1, Tagln, Myl9, Sparc*) and the kidney-fibroblast-enriched gene *Pax8;* Kidney2 strongly expressed epithelial genes (*Krt7, 8, 18*, *Epcam, Clu*) (**Supplementary Fig. 12 d-e, table 6**). As for the cultured heart fibroblasts, Heart1 displayed myofibroblast genes (*Acta2*, *Tagln, Myl9)* and Heart2 had enhanced signature of injury response/acutely activated fibroblasts (*Mt1, Ccl2, Clu, Dcn)* (**Supplementary Fig. 12 a-b, Table 6**) [22]. Overall, scRNAseq experiments showed that cultured cells present an activated/myofibroblast-like phenotype compared to freshly isolated cells and confirmed the retention of an organ-specific core transcriptome identity.

### Organ enriched transcriptome is involved in the fibrotic response

To investigate the functional relevance of organ-enriched fibroblast transcriptomes, a CRISPR knock-down approach was used to down-regulate core organ transcription factors, taking the heart as a model (**Fig. 5**). *Rosa^Cas9-GFP^* [36] adult cardiac fibroblasts were co-transfected with mCherry mRNA and GFP guide RNAs for determination of transfection and knock-down (KD) efficiency, respectively (**Fig. 5a**). After 72h of transfection, mCherry was observed in roughly 67% of cells (**Fig. 5b**) by flow cytometry, and GFP mRNA expression was downregulated by over 90% when compared with scrambled guides (negative control) (**Fig. 5c**). GFP fluorescence was also dramatically decreased by 72 hours (**Fig. 5a**) [37].

**Fig. 5:**
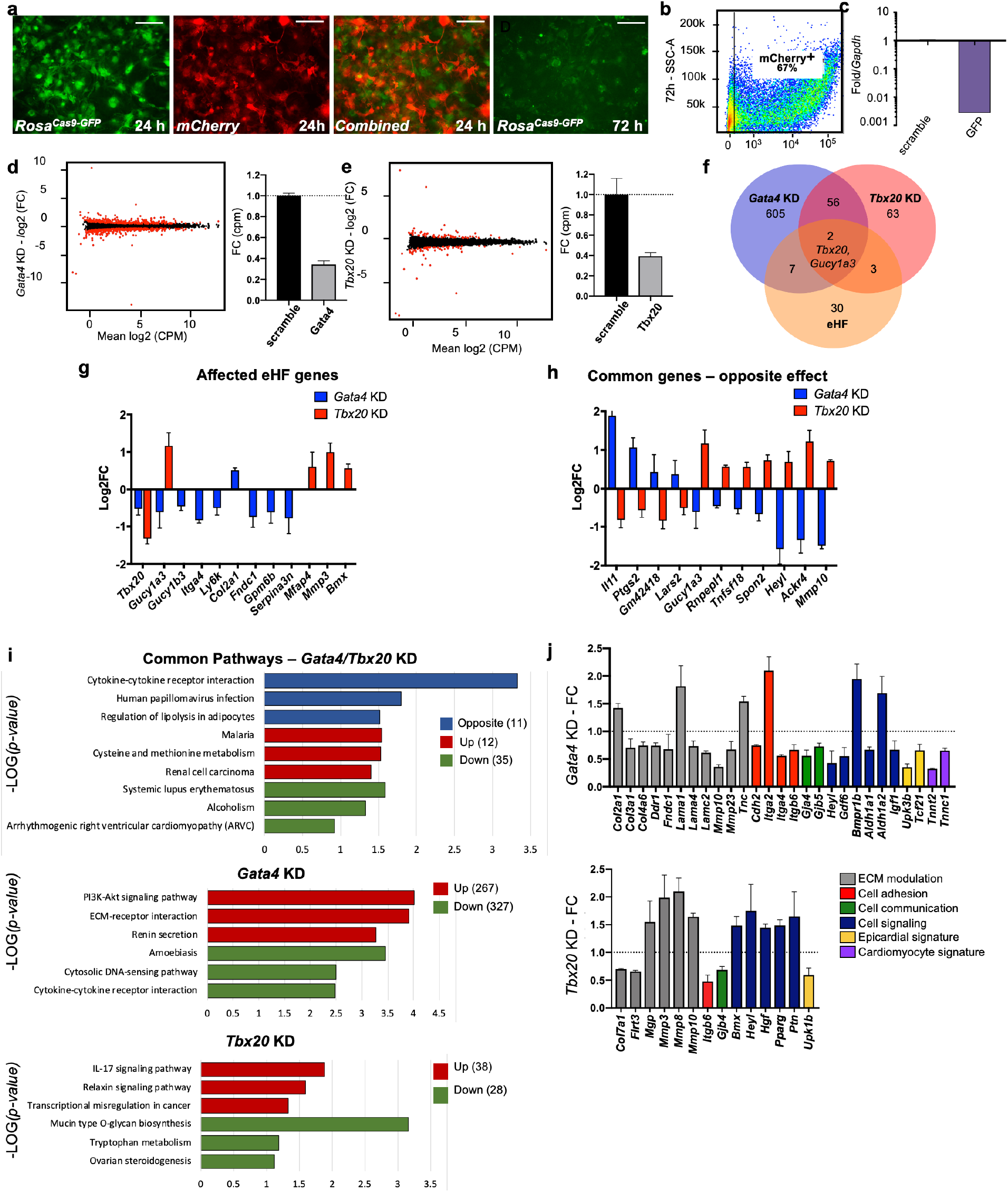
In vitro knock-down of core cardiac transcription factors. **a** Images of adult cardiac fibroblasts derived from *RosaCas9-GFP* mice, 24 and 72 hours post-transfection with 2 guide RNAs for *GFP* and CleanCap® mCherry mRNA. **b** Flow cytometry plot showing the expression on mCherry in 67% of transfected cells (representative image of 3 independent experiments). **c** Relative quantification qPCR of *GFP*, normalized by *Gapdh* expression, in cells transfected with scrambled or GFP guide RNAs. **d** Volcano plots showing genes differentially expressed in *Gata4* (left) and *Tbx20* knock-downs (right). Fold change in expression of *Gata4* (left) and *Tbx20* (bottom) in cells transfected with specific gRNAs versus scramble RNA, quantified through RNA sequencing. **f** Venn diagram showing the overlap among genes affected by Gata4 (*Gata4KD*) or Tbx20 (*Tbx20KD*) knock-down and number of genes upregulated by 10-fold or more in heart fibroblasts (eHF) compared to other organs**. g** plot showing changes in expression of eHF genes affected by *Gata4* (in blue) or *Tbx20* (in red) knock-down**. h** Genes regulated by both *Gata4* and *Tbx20* in opposing manner. **i** KEGG Pathway analyses. Top panel: 58 genes affected in both *Tbx20* and *Gata4* knock-down; Middle panel: 594 genes affected by *Gata4* knock-down; Bottom panel: 66 genes affected by *Tbx20* knock-down. Blue - pathway changed in opposite directions, Red - up-regulated pathways, Green – down-regulated pathways. **j.** Hand-picked genes illustrate alterations in processes known to affect the cardiac fibrotic response for *Gata4* or *Tbx20* knock-down. All data are represented as fold changes over scrambled control (Average ± SEM; **d-j**) from bulk RNA-seq of 3 biological replicates per condition.

With this confirmation, *Gata4* and *Tbx20*, core transcription factors essential for heart formation in embryonic development [38], were knocked-down in cultured *Rosa^Cas9-GFP^* cardiac fibroblasts, followed by bulk RNA-seq analysis (**Fig. 5d-j**). *Gata4* is expressed by all organ fibroblasts while fibroblast *Tbx20* expression is restricted to the heart. Despite similar KD efficiencies for both targets (∼60%; **Fig. 5d-e**), *Gata4KD* induced a higher number of dysregulated genes (red dots on volcano plots) compared with *Tbx20KD*. Among genes up-regulated by 10-fold in the original bulk organ analysis (eHF; **Fig. 5f-g**), 9 were dysregulated by *Gata4KD* and 5 by *Tbx20KD.* Only 2 of these genes were dysregulated in both conditions, including *Tbx20*, suggesting *Gata4* as a possible upstream regulator of *Tbx20*. A number of genes showed opposite regulation between *Gata4KD* and *Tbx20KD* (**Fig. 5h**), confirming the specificity of KD response. These included cytokines and cytokine-receptors (*Il11, Tnfsf18, Ackr4*), genes involved in infection (*Ptgs2, Heyl*), cell adhesion and migration (*Spon2*, *Mmp10*). Of note, among the genes selectively upregulated in *Gata4KD*, *Il11* is a key mediator of organ fibrosis, possible downstream effector of TGF β [39]. Among genes upregulated in *Tbx20KD, Heyl*, downstream effector of Notch, is involved in cardiogenesis and is thought to repress *Gata4* expression [40], *Mmp10* is upregulated in patients with end-stage HF [41], and it is involved in valve ossification [42]. KEGG pathway analyses (**Fig. 5i, Supplementary table 7**) confirmed the involvement of *Gata4* and *Tbx20* in common but also diverse pathways. The top pathways uniquely up-regulated by *Gata4KD* included Akt signaling, ECM-receptor interaction and renin secretion, implicating *Gata4* in the modulation of cardiac fibroblast growth and ECM changes [43–45]. The top pathways up-regulated by *Tbx20KD* involved IL-17 and relaxin signaling, as well as transcription misregulation in cancer. IL17 has been shown to regulate the fibrotic response in pro-inflammatory conditions such as psoriasis and pulmonary/liver fibrosis [46–49], while relaxin has a well-established role in suppressing myofibroblast activation and ECM remodeling [50–52].

In summary, both gene KDs affected matrix components and modulators (**Fig. 5j**), as well as cell adhesion, cell-cell communication and cell signaling genes. Markers of the epicardium, the external layer of the heart from which embryonic fibroblasts derive [53], were also modulated, as well as several myocardial genes, found in low levels in cardiac fibroblasts and downregulated in *Gata4KD*. These results confirm the biological relevance of organ-specific fibroblast gene expression.

### Organ fibroblast specificity affects tissue function in co-culture systems

The studies described above confirmed that fibroblasts retain an organ-specific transcriptome from embryonic development to adulthood, and that their identities are largely maintained in cultured cells, suggesting that fibroblast transcriptomes may be important for *in vivo* organ function. To determine whether the source of organ fibroblast affects organ function, 2D and 3D co-cultures of cardiomyocytes (CMs) with adult kidney and cardiac fibroblasts were performed (**Fig. 6**).

**Fig. 6:**
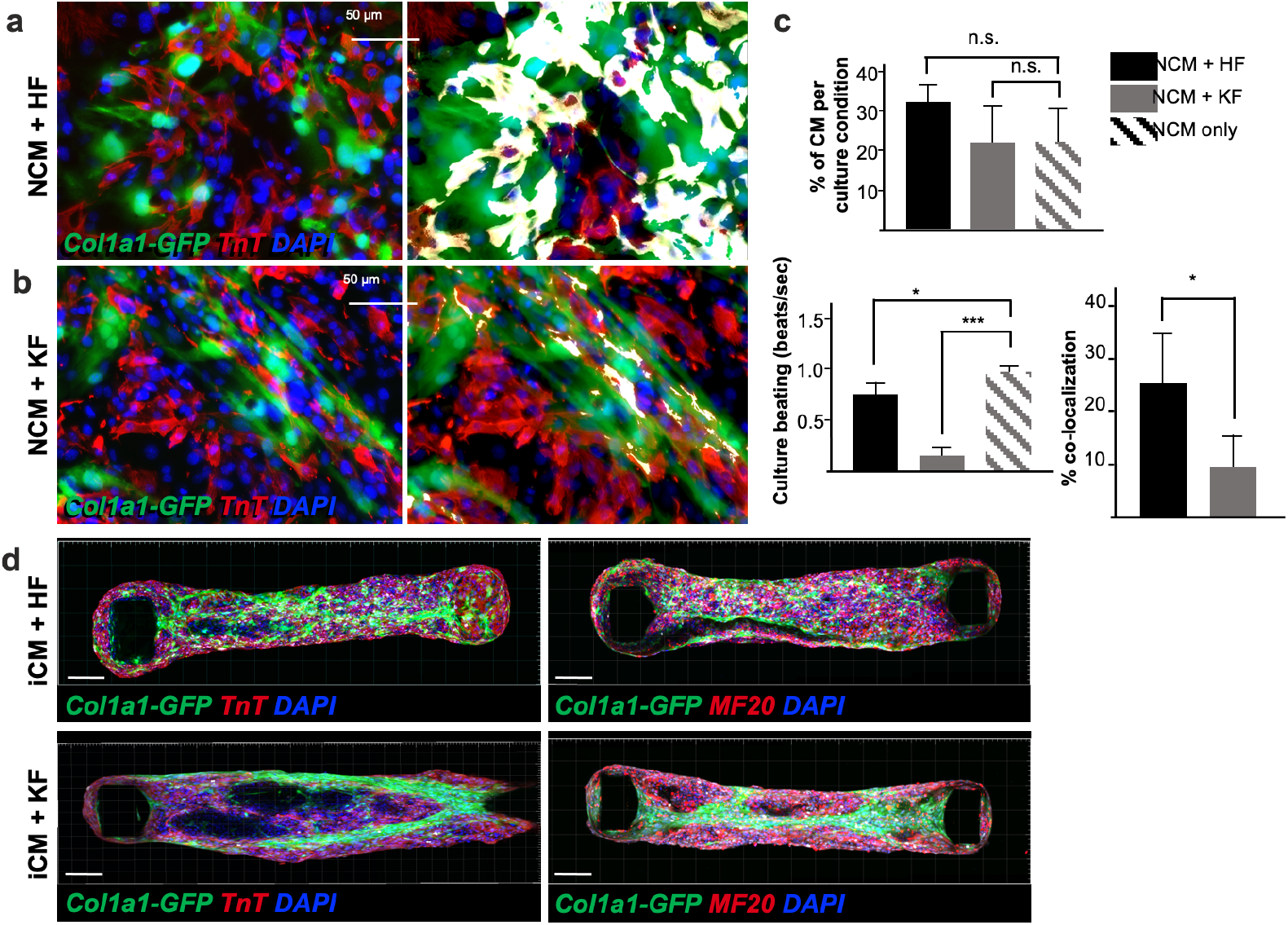
Adult fibroblasts retain tissue specific function *in vitro*: impaired neonatal cardiomyocyte beating in presence of kidney derived fibroblasts. **a-b** Immunocytochemistry for TnT on 2D co-culture of neonatal ventricular cardiomyocytes (NCM, TnT+ in red) with either adult cardiac fibroblasts (HF) in **a** or kidney fibroblasts (KF) in **b,** isolated from *Col1a1-eGFP* mice (in green). Nuclei are labelled with DAPI (in blue). The right panel shows colocalization of the two cell types (green+ and red+) in white. **c** quantifications of the percentage of cardiomyocytes per culture condition (top), beating of the 2D cultures expressed in beats per second (bottom left) and percentage of co-localization (bottom right). **d** Confocal Z-stack images reconstructed with Imaris, of cardiac microtissues constituted of 85% hiPSCs derived Cardiomyocytes (iCM) and 15% of either cardiac (HF) or kidney (KF) fibroblasts, stained for TnT (in red, left panels) or MF20 (in red right panels). The adult fibroblasts were isolated from *Col1a1-eGFP* mice (in green), nuclei stained with DAPI (in red). Scale bar= 50μm. All data in (**c**) are mean ± SEM on 3 independent experiments, p-values were calculated by two-sample t-test.

For 2D cultures, neonatal ventricular CMs were plated with *Col1a1-GFP*+ fibroblasts from adult kidney or heart [54]. Within 24h, coculture with adult kidney fibroblasts almost completely impaired CM contractility (**Fig. 6b-c, Supplementary Video 1)**, although the number of CMs present in both cultures was not significantly different. Conversely, cardiac fibroblast co-culture resulted in a syncytium of cells beating in synchronism (**Supplementary Video 2**) at a relatively lower pace than neonatal CMs alone (**Fig. 6c**), possibly reflecting an effect of adult HF on neonatal CM maturation [55]. In addition, cardiac fibroblasts were well integrated with neonatal CMs, as shown by the percentage of co-localization, while kidney fibroblasts and CMs seemed to repel each other. To confirm these findings, 3D cardiac microtissues we generated, as previously described [56, 57]. A suspension of 85% human induced pluripotent stem cell derived CMs (iCMs) and 15% adult cardiac or kidney fibroblasts was loaded on millitissue devices with pairs of cantilevers to generate force. As expected, cardiac fibroblasts were homogenously interspersed in the organoids, while kidney fibroblasts were aggregated to the center or periphery of the organoids (**Fig. 6d**), indicating lack of integration between the two cell types. These results implicate organ-specific fibroblasts in imparting their cognate tissue integrity.

### Ectopically transplanted fibroblasts retain core transcriptional identity

To investigate if adult fibroblasts maintain their organ-specific signature when exposed to a different tissue microenvironment *in vivo*, fibroblasts from tail, heart and kidney of *ROSA^mT/mG^* mice were absorbed on surgical gel foam and transplanted under the kidney capsule of syngeneic C57BL6/J mice (**Fig. 7a**). Three days post-transplantation, kidneys were dissected and sorted (**Fig. 7b**) to determine transcriptional changes in transplanted cells. Transplanted heart (HFs), kidney (KFs), tail (TFs) fibroblasts, and corresponding *in vitro* cultured controls (HFc, KFc, TFc) were processed for bulk RNA-seq. Multidimensional scaling plot for all samples showed that the three organ fibroblast types retained a distinct identity post-transplant, despite a reduced transcriptomic separation (**Fig. 7c**).

**Fig. 7:**
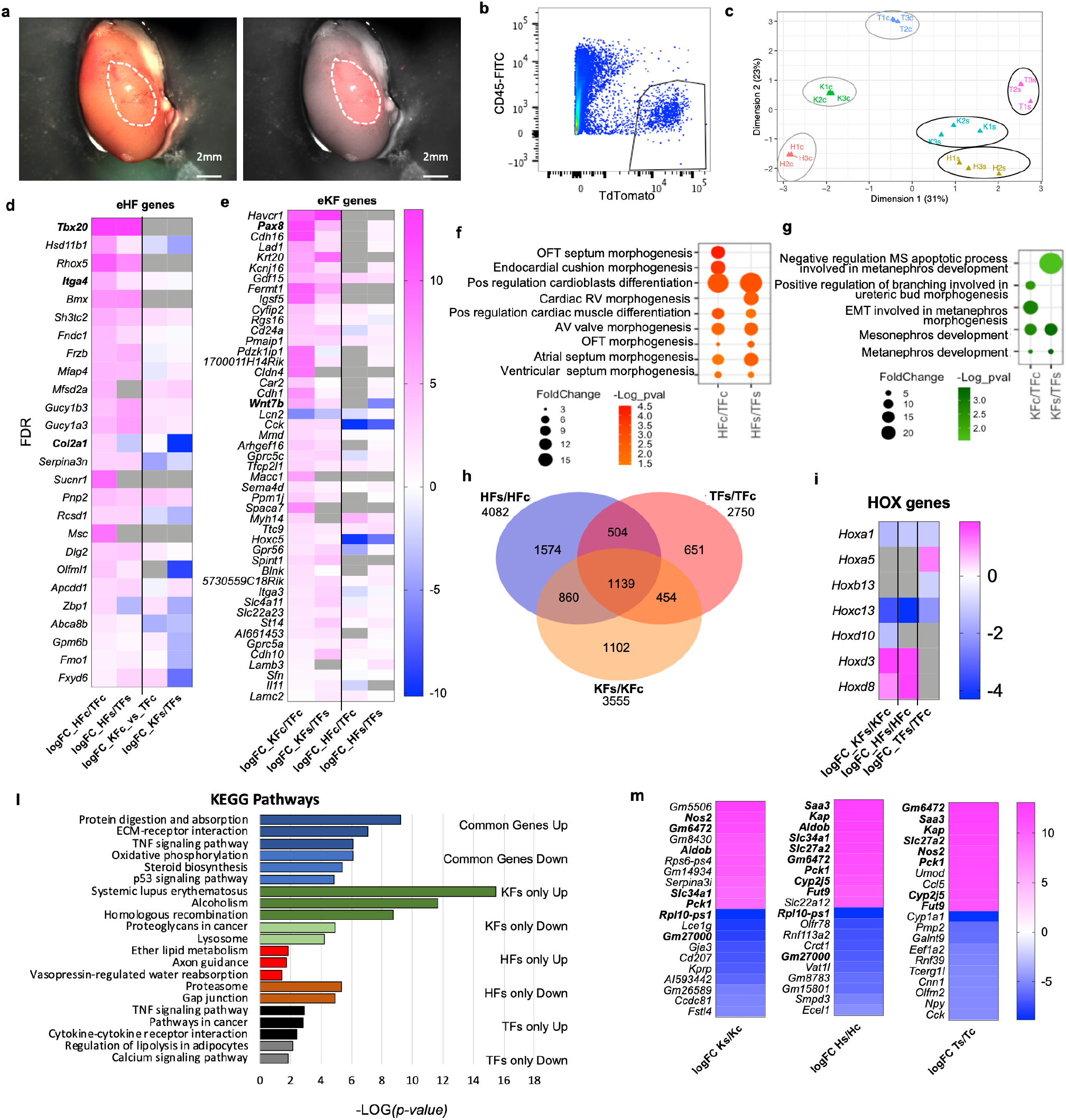
Fibroblasts’ tissue-specific response to *in vivo* ectopic transplantation under the kidney capsule. **a** Representative images of a dissected kidney: brightfield image on the left; brightfield overlaid with the fluorescence image acquired in the red channel on the right, to highlight the area where TdTomato+ cells were transplanted. **b** Flow cytometry plot showing the gating strategy used to isolate live CD45-TdTomato+ cells 3 days post-transplantation in the kidney capsule. **c** Multidimensional scaling plot calculated on the top 500 genes post-normalization, to visualize the level of transcriptomic similarity among all the samples. **d-h** analysis of the cell-specific identity by comparing the HF and KF gene expression to TF in culture (HFc/TFc, KFc/KFc) and post-transplant in the kidney capsule (HFs/TFs, KFs/KFs). (**d**) Heatmap showing the expression of significantly regulated eHF genes in the 4 conditions. (**e**), Heatmap showing the expression of eKF genes in the 4 conditions. (**f**) dot plot indicating the GO terms associated to cardiac development, identified from the DAVID database analysis of HFs or HFc enriched genes, (**g**) dot plot indicating the GO terms associated to cardiac development, identified from the DAVID database analysis of KFs or KFc enriched genes**. h-m** analysis of the differential expression by experimental condition: transplanted cells versus cells in culture HFs/HFc, TFs/TFc, KFs/KFc. (**h**) Venn diagram showing the significant differentially regulated genes (FDR<0.05) in the three comparison sets. Only genes with a logFC>1 or <-1 were considered. (**i)** Heatmap of differentially regulated Hox genes. (**l**) Bar plot of the KEGG pathway analysis on the common regulated genes (light blue-downregulated, dark blue-upregulated), and genes uniquely modulated in KF (dark green-upregulated, light green-downregulated), HF (red-upregulated, brown-downregulated), TF (black-upregulated, grey-downregulated). **(m)** Heatmaps showing the top 10 upregulated and top 10 downregulated genes for each dataset. In bold are the genes shared in two different sets of comparisons. Data are means of 3 biological replicates per each condition. All the heatmaps **d-e-l-m** show the average logarithmic fold change. For **d-e** genes were ordered based on the FDR (smaller to larger value) of the comparison in the first column, HFc/TFc and KFc/TFc respectively. Dotplots **f-g** and bar plots **h-m** data were organized based on the logarithmic transformation of the p-values (-log(p-value)). eKF = kidney fibroblasts enriched genes, same as shown in Suppl. Fig.8, eHF= heart fibroblasts enriched genes, same as shown in Suppl. Fig.10, KFs= transplanted kidney fibroblasts, HFs=transplanted heart fibroblasts, TFs=transplanted tail fibroblasts, KFc= kidney fibroblasts in culture, HFc=heart fibroblasts in culture, TFc= tail fibroblasts in culture

To assess eventual changes in organ-specific identity, we compared the expression of sorted and cultured heart and kidney fibroblasts with the correspondent tail fibroblasts, and we analyzed the expression of heart-enriched (eHF) and kidney-enriched (eKF) genes identified from the initial bulk RNA analyses (**Supplementary Table 3,** **Figs. 5-10**). We observed that fibroblasts generally maintained their core identity after transplant. Of the 26 genes enriched in HFc, 21 (80.7%) were similarly modulated in HFs, only 2 were downregulated and 3 were not detected (**Fig. 7d**). As expected, eHF gene expression was low or downregulated in KFc compared to TFc and kept a similar expression pattern post-transplant (**Fig. 7d**). Of the 47 eKF significantly expressed in KFc, 41 (87.2%) were modulated in the same direction in KFs, one gene was downregulated, and 5 genes were not detected. 26 (55.3%) eKF genes were also found in HFc, 17 mildly upregulated, 9 downregulated. Of these, 14 were similarly regulated in HFs, one gene was not detected and 11 were differentially regulated (7 upregulated, 4 downregulated). An additional 13 eKF genes were detected in HFs, all mildly upregulated except for 1 downregulated, showing an adaptation to the new microenvironment (**Fig. 7e**). Gene ontology (GO) analysis of KF or HF enriched genes in culture or post-transplant using *DAVID* (Database for Annotation, Visualization and Integrated Discovery [58]) revealed terms related to organ development, in line with the previous observations (**Fig. 7f-g**). The top GO terms for both HFc and HFs were related to cardiac morphogenesis and cardioblast differentiation; the top terms for KFc and KFs were related to mesonephros and metanephros development. In summary, both HFs and KFs maintained their core transcriptomic identity compared to TFs.

### Ectopically transplanted fibroblasts adapt to a new microenvironment

To analyze the differential HF, KF, TF responses to the transplantation, gene expression of each post-transplant fibroblast type was compared with its equivalent control kept in culture. Despite the retention of organ identity signatures, numerous genes were modulated in transplanted fibroblasts compared to cultured controls (4082 genes for HFs/HFc, 3555 for KFs/KFc, and 2750 for TFs/TFc) (**Fig. 7h**). These included 4-5 Hox genes per cell type, showing a modulation of the cell-type specific positional code. *Hoxa1* (the highest expressed in HFc) and *Hoxc13* were downregulated in all conditions (**Fig. 7i**). The tail-enriched *Hoxb13* was downregulated in TFs, while *Hoxa5* was increased. *Hoxd3* and *Hoxd8* were upregulated in both HFs and KFs; and *Hoxd10* was downregulated in KFs. Interestingly, *Hoxd8* is important for the maintenance of epithelial phenotype in adult kidney and is expressed in the ureteric bud during development [59], while *Hoxd10* is diffusely expressed in kidney mesenchyme in embryos; both *Hoxd10* and *Hoxd3* regulate *Itga3* expression and have been involved in different types of cancer [60]. The differential regulation of *Hoxd10* and *Hoxd3* may suggest the acquisition of a cortex-like phenotype by transplanted KFs.

KEGG analysis of the genes modulated in response to the transplant showed upregulation of pathways related to fibrosis and damage response (TNF signaling, ECM-receptor interaction, protein digestion and absorption) and downregulation of oxidative phosphorylation, steroid biosynthesis, and p53 signaling among the commonly regulated genes (**Fig. 7l, Supplementary Fig. 13a**). Genes uniquely upregulated in KFs/KFc were related to homologous recombination and cell cycle, with pathways including several histone genes (Alcoholism, Systemic lupus erythematosus, **Supplementary Fig. 13a)**; those selectively upregulated in HFs/HFc were associated with cell migration (axon guidance), vasopressin regulated water absorption and lipid metabolism; and to pro-inflammatory pathways for TFs/TFc (TNF signaling, cytokine-receptor interactions, pathways in cancers) (**Fig. 7l**). Similarly, IPA analysis revealed that the most significantly affected Canonical Pathways were related to fibrosis (Hepatic fibrosis signaling, GP6 Signaling); cell migration (Axonal Guidance Signaling); acute phase response, inflammation and cholesterol biosynthesis (**Supplementary Fig. 13b**). Interestingly, Cardiac Hypertrophy Signaling (which include many pro-fibrotic signals like AngII, TGFb, IGF1), and HIF1a signaling were predicted to be downregulated in KFs and upregulated in HFs and TFs, possibly inferring a better resilience of KFs to the kidney capsule environment.

Among the top 10 upregulated genes in transplanted fibroblasts, 6 were shared by the ectopically transplanted HFs and TFs, including the serum amyloid A *Saa3,* secreted during the acute phase of inflammation[22]; two metabolic enzymes, *Slc27a2,* primarily expressed in kidney and liver involved in lipid biosynthesis and fatty acid degradation, and *Pck1* key regulator of gluconeogenesis; the cytochrome gene *Cy2j5* involved in vasorelaxation[61]; the kidney abundant protein *Kap,* androgen-regulated, proximal tubule–specific not expressed at detectable levels in tissue other than the kidney[62]*, Fut9* a fucosyltransferase with the highest expression in adult pancreas, placenta, kidney (**Fig. 7m**). In summary, while transplanted fibroblasts maintained their core identity, they responded to the kidney microenvironment by expressing a subset of kidney-specific genes, modulating positional code genes (i.e. *Hoxa1* for heart and *Hoxc13* for tail), and activating common and cell-specific pathways in the attempt to adapt to the new, more hypoxic condition.

## Discussion

The ability to target specific organ fibroblasts has long been impaired by the misconception that fibroblasts were functionally and phenotypically homogeneous cells, deputized to synthetizing and organizing the extracellular matrix, an idea possibly fostered by a common embryonic origin in the primary mesenchyme [13]. Indeed, a recent study has shown that, despite apparent tissue-specific imprinting, fibroblasts subclusters across multiple organs present a common hierarchy, with two universal subtypes generating more specialized or activated fibroblasts states [63]. However, recent advances in lineage tracing and single-cell transcriptomic have revealed an extensive intra- and inter-organ heterogeneity [3, 13]. Multi-organ studies show that B-cells [64], endothelial cells [65], fibroblasts [21, 30] transcriptomes tend to cluster separately based on the organ of origin, thus putting into question the very concept of cell type. For fibroblasts isolated from muscular tissues, the organ-specific differences are considered to be mostly attributable to changes in the matrisome [21].

We previously reported that fibroblasts isolated from the adult mouse heart retain a cardiogenic transcriptional program [23]. Here, we compared primary cultures of fibroblasts isolated from organs of different anatomical positions to expand our previous analysis and assess whether development-related genes contribute to the fibroblasts’ inter-organ functional heterogeneity. The results of this analysis highlight the presence of an organ-enriched positional code, and the expression of core genes that represent the developmental signature of fibroblast organ origin previously thought to be restricted to the parenchymal component. These molecular profiles are established during embryogenesis, reflecting the fact that organ fibroblasts are not generated from a common progenitor pool, but arise independently in different body segments and organs during embryonic development and persist to adulthood.

As in our previous study [23], we chose to analyze cultured fibroblasts to reduce the risk of contamination from parenchymal cell mRNA. Fibroblast expression patterns in culture were recapitulated in freshly isolated single cells, mostly enriched in activated or myofibroblast-like fibroblast subclusters. These gene signatures can predict the tissue of origin of a mixed population of primary cultured cells analyzed at the single-cell level. Using the heart as an example we show that signature genes contribute to organ fibroblast function, as evidenced by the deregulation of several pro-fibrotic and pro-inflammatory genes with knock-down of core transcription factors *Gata4* (expressed in all fibroblasts types) and *Tbx20* (cardiac-specific) in cultured adult cardiac fibroblasts. These results place *Gata4* upstream of *Tbx20,* both of which upregulate distinct pro-fibrotic signals, modulate genes involved in extra-cellular modulation and cell adhesion, and have opposite effects on cytokine-cytokine receptor expression, confirming that the core cardiogenic program in cardiac fibroblasts is involved in regulating their function.

Dermal fibroblasts from different sites of the body have shown different efficiency of reprogramming into induced pluripotent stem cells [66], but not much is known about other fibroblast tissue-specific functions. The co-culture studies presented here further reinforce the importance of fibroblast core transcriptomes for specialized organ function: while interspersion of cardiac fibroblasts within CM cultures facilitated the propagation of the electric pulse forming a syncytium, co-cultured kidney fibroblasts clustered separately and inhibited CM contraction, both in 2D and 3D assays. These findings carry repercussions to *in silico* organ bioengineering, where combining the correct match of diverse organ cell types may be essential for proper organ formation. Indeed, human induced pluripotent stem cell derived cardiac stromal cells enhance maturation of cardiac microtissues [67]. In addition, the source and type of organ scaffolding, mainly deployed by fibroblasts, is essential for the re-creation of organs in a dish [68].

Previous studies have shown that skin fibroblasts and mesenchymal cells from different organs keep a positional identity [17–19]. For mesenchymal cells, the Hox code was maintained also in culture, although whether this depended on cell-to-cell contact remained to be determined [19]. Here we show that adult fibroblasts, isolated from a variety of organs, preserve the expression of Hox genes in culture, but that tissue-specific Hox genes were downregulated after ectopic transplantation under the kidney capsule, suggesting that cellular environment can induce reprogramming of positional codes. Interestingly, control kidney fibroblasts also presented changes in the Hox code after transplantation, possibly reflecting the adaptation to the space between the capsule and the cortex, with the decrease of the mesenchyme gene *Hoxd10* and increase of *Hoxd3* important for the maintenance of an epithelial phenotype in adult kidney. All three transplanted fibroblast types (heart, tail, kidney) presented an activated phenotype, involving initiation of the acute phase response, pro-fibrotic signals, and metabolic changes. While transplanted heart and tail fibroblasts showed a clearer activation of pro-inflammatory pathways, genes associated with cell migration, and HIF1a signaling; transplanted kidney fibroblasts appeared more resilient in their native milieu, with the activation of pathways related to proliferation and upregulation of genes indicative of a more epithelial cortex-like phenotype. Both heart and kidney fibroblasts retained a “memory” of their organ of origin, defined by the resilient expression of the core of development-related genes when compared to tail fibroblast control. It remains to be determined if this memory can be erased by a longer residence in the ectopic microenvironment. In light of recent studies on endothelial cells [69, 70], we propose that the expression of organ-specific genes, previously thought to be restricted to parenchymal cells, may form the basis for organ cohesiveness and performance.

In summary, adult fibroblasts maintain a lasting blueprint of the organ in which they reside, reflective of its developmental origin, which likely plays a role in the orchestration of the tissue-specific homeostasis and reparative response. Exploiting the organ-specific properties of fibroblasts may be a valuable strategy for the targeted control of organ fibrosis, an integral feature of organ failure and disease progression affecting a multitude of pathologies.

## Methods

### Mice

All experiments were performed with young adult (8-12 weeks old) C57BL/6J, Gt(ROSA)26Sor ^tm1.1(CAG-cas9*,-EGFP)Fezh/^J (Rosa^Cas9-EGFP^), Rosa26^mt/mg^ (JAX Stock# 007576)[71], Col1a1-GFP[72] male mice and Col1a1-GFP embryos E16.5. All animal experimentation conformed with local (Jackson Laboratory) and national (NHMRC and NIH) guidelines, under IACUC protocol 16010.

### Fibroblast isolation and sorting

Liver, heart, lung, kidney, tail, gonad and ventral skin of adult mice and E16.5 embryos were dissected and finely minced. Fibroblasts were isolated using enzymatic digestion with 0.05% Trypsin/EDTA (Gibco) under agitation at 37°C for 40 minutes. Cells were spun and plated in 10cm dishes and cultured to semi-confluence in DMEM (ThermoFisher) high glucose supplemented with 10% FBS (ThermoFisher), sodium pyruvate and pen/strep (ThermoFisher) in a 5% CO_2_ incubator at 37°C. Passage 0 cells were then trypsinized using TriplE (ThermoFisher) and further processed for flow cytometry, labeled using CD90-AF647 (BioLegend), CD45-PeCy7 and CD31-Pe (eBioscience) in 2%FBS/HBSS (ThermoFisher) and sorted using Influx or Aria II Sorter (BD). The CD90+; CD45-; CD31-fraction was collected for mRNA isolation (**Supplementary Fig.1**). Adult fibroblasts from Rosa^Cas9-EGFP^ and Col1a1-GFP were sorted using CD90-APC (BioLegend), CD45-APCCy7(BioLegend), CD31-PECy7 (BD) after 3 or 5 days respectively.

### Microarray Assay

Sorted organ fibroblasts were resuspended in cell lysis buffer, further processed for total RNA isolation using the RNAqueous Micro kit (ThermoFisher) and DNAse digested on column. Fibroblasts from individual mice were used for each replicate. Triplicates or more were used for each organ. Samples were further processed by the Monash Health Translational Precinct Medical Genomics Facility and ran on Agilent SurePrint G3 mouse gene expression arrays (single color).

### Bulk RNA sequencing

Total RNA was isolated from heart tissue using miRNeasy Mini kit (Qiagen); from cultured fibroblasts (CRISPR-experiment) and sorted fibroblasts (kidney capsule experiment), using RNeasy Micro kit (Qiagen) according to manufacturer instruction and including the optional DNase digest step. Sample concentration and quality were assessed using the Nanodrop 2000 spectrophotometer (Thermo Scientific) and the Total RNA Nano or Pico assays (Agilent Technologies).

For human heart samples, libraries were constructed using the KAPA RNA Hyper Prep Kit with RiboErase (HMR) (KAPA Biosystems), according to the manufacturer’s instructions. For cultured fibroblasts (CRISPR-experiment), libraries were constructed using the KAPA mRNA HyperPrep Kit (KAPA Biosystems), selecting polyA containing mRNA using oligo-dT magnetic beads, according to the manufacturer’s instructions. For cells isolated from the kidney capsule, given the low RNA input, libraries were constructed using the SMARTer Stranded Total RNA-Seq Kit v2-Pico (Takara), according to the manufacturer’s protocol.

All the libraries were checked for quality and concentration using the D5000 ScreenTape assay (Agilent Technologies) and quantitative PCR (KAPA Biosystems), according to the manufacturers’ instructions; pooled and sequenced 75 bp paired-ended (human samples) or single-end (cultured and sorted fibroblasts) on the NextSeq 500 (Illumina).

### Single cell RNA sequencing

Fibroblasts isolated from the different tissues were FAC-sorted and loaded onto a single channel of the 10X Genomics Chromium single cell platform. Briefly, cells were loaded for capture using the v2 single cell reagent kit. Following capture and lysis, cDNA was synthesized and amplified (14 cycles) as per manufacturer’s protocol (10X Genomics). The amplified cDNA was used to construct an Illumina sequencing library and sequenced on a single lane of a HiSeq 4000.

### Bioinformatics Analyses

For microarray experiments, data extraction and pre-processing were performed as described previously [73]. In brief, raw single-channel signals were extracted (Agilent Feature Extraction Software v.11.0.1.1), and quality control was performed using the default “Compromised” option in (GeneSpring GX v.12.6), with threshold raw signal of 1.0. The approximate mean of 24 samples × ∼55,000 probes (10,000) was used as a natural threshold between high-intensity probes and low-intensity probes. If several probes represented a single gene, the mean of these probes was used. Probes that could not be mapped to any gene were discarded. Log-2 transformation and quantile normalization was done using R scripts and public Bioconductor packages. Differential analysis was performed using the Bioconductor *limma* package, which fits a linear model to the gene expression data, revealing the differential expression patterns (Benjamini-Hochberg adjusted p-value<0.05 and fold-change >2). These genes were extracted from the transcriptome to generate a heat-map together with hierarchical clustering dendrograms using MultiExperiment Viewer (MeV) [74]. Differentially expressed genes showing more than 10-fold change in any given organ were retrieved and an interaction file listing in which organs these genes were enriched was constructed. The interaction file was used as input for Cytoscape [75] in order to reconstruct the network of genes shared by two or more organs, or only specifically enriched in one organ. The network layout was constructed using a Spring Embeded layout and MeV. Gene Ontology over-representations for the organ-specific subset of genes was performed using the Cytoscape Bingo plug-in. Ingenuity Pathway Analysis was performed using the IPA software (Qiagen).

For single cell RNA sequencing analyses of freshly isolated fibroblasts, stromal cell data from the Mouse Cell Atlas [30] were kindly provided by Dr. Guoji Guo and Dr. Huiyu Sun. The data were re-analyzed using Seurat v3 [76]. Cell with less than 200 and more than 2500 transcripts were filtered out. Out of the original aggregate, containing 21 samples and 4830 cells, 5 populations of interested were selected for further analysis: “Lung”, “Testis”, “Kidney”, “Liver”, “NeonatalHeart”, corresponding to 682 cells. Data were natural-log normalized and scaled using the top-2000 most variables features in the raw data. Principal component analysis (PCA) dimensionality reduction was calculated on 50 principal components; the Uniform Manifold Approximation and Projection (UMAP) dimensional reduction was calculated on 24 dimensions; cluster determination was performed using shared nearest neighbor (SNN) at a 0.5 resolution. Cluster’s markers genes were identified with the *FindAllMarkers* function, using the default Wilcoxon Rank Sum test, at a threshold of 0.25 and a minimum difference in the fraction of detection (*min.diff.pct*) of 0.3. Pairwise comparisons were done using the *FindMarkers* function, with MAST assay and only testing genes that are detected in 25% of cells in either of the two populations (*min.pct*=0.25).

For bulk RNAseq analysis on cultured fibroblasts post-CRISPR-Cas9 knock down or kidney capsule implant: Single end, Illumina-sequenced stranded RNA-Seq reads were filtered and trimmed for quality scores > 30 using a custom python script. The filtered reads were aligned to Mus musculus GRCm38 using RSEM (v1.2.12) which performed alignment using Bowtie2 (v2.2.0) (command: rsem-calculate-expression -p 12 --phred33-quals --seed-length 25 --forward-prob 0 --time --output-genome-bam -- bowtie2). RSEM calculates expected counts and transcript per million (TPM). The expected counts values from RSEM were used in the edgeR 3.20.9 package to determine differentially expressed (DE) genes (based on fold-change > 1 and FDR < 0.05).

For single cell RNA sequencing data from cultured fibroblasts, Illumina basecall files (*.bcl) were converted to FASTQ files using Cell Ranger v1.3, using the command-line tool *bcl2fastq* v2.17.1.14. FASTQ files were then aligned to mm10 genome and transcriptome using the Cell Ranger v1.3 pipeline, which generates a gene vs cell expression matrix. The data were analyzed using Seurat v3 [76] using the same pipeline and parameters as described above, unless stated below. Given the high average number of features, cell with less than 200 and more than 8500 transcripts were filtered out, obtaining 1121 cells. Data were normalized and scaled as described above. PCA dimensionality reduction was calculated on 50 principal components; UMAP dimensional reduction was calculated on 28 dimensions (value chosen based on the *ElbowPlot* of the standard deviations of the principal components).

### qPCR

cDNA synthesis of RNAs used for the microarray was performed using the Superscript VILO kit (Invitrogen) following manufacturer’s instructions. PCR reactions were performed using GoTaq Green master mix (Promega). qPCR reactions were performed using SYBR green master mix (Roche) and analyzed using the LightCycler 480 (Roche). At least 2 individual experiments in triplicate were performed. We tested several primers for endogenous control (*Tbp*, *Gapdh*, *L13*, *Ppi*, *Actab* and *Hprt*) and chose *Hprt* for further experiments due to its consistent reproducibility within and among samples (**Supplementary Fig. 1**). Primers are described in **Supplementary Table 8**. All PCR reactions were performed in triplicates and repeated at least twice per sample. Standard error of the mean is represented in all graphs. Prism v7.0 was used for the generation of graphs and statistics.

### Neonatal mouse cardiomyocyte isolation

The protocol from neonatal cardiomyocytes isolation was adapted from Argentin S. et al[77]. Hearts were collected from litters of 1-3days old pups, cut open and transferred to trypsin (1mg/ml in HBBS pH6.4) for overnight digestion at 4°C. The next day, hearts were subjected to 3 x 5min digestions with Collagenase II (1mg/10ml; Worthington). The cell suspension was collected in DMEM containing 10% fetal calf serum (FCS) and passed through a 100um cell strainer. After 5min centrifugation at 1000rpm, cells were plated in 10cm dishes. Two rounds 1-hour pre-plating were done to remove cells highly adherent to plastic such as fibroblasts, before seeding the cell suspension on plates coated with 1:200 fibronectin (ThermoFisher) in 0.1%gelatin (ThermoFisher).

### 2D Co-cultures

Adult fibroblasts isolated from heart or kidney of Col1a1-GFP mice were cultured to semi-confluence for 3-5-days, after which they were resuspended and co-cultured with mouse neonatal cardiomyocytes at a 4:1 ratio. After 24h, media was changed to DMEM containing 2% FCS, videos were recorded, and cells were imaged with an Eclipse Ts2 inverted fluorescence microscope (Zeiss) and fixed with 4%PFA for 10min at 4°C for further staining.

### Cardiac Microtissues

Cardiac Microtissues were generated as previously described[56, 57], using polydimethylsiloxane (PDMS) 3D microarrays with 24 microwells containing cantilevers. A suspension of 1.3 million cells, 85% hiPSCs derived Cardiomyocytes (iCM) and 15% cardiac or kidney fibroblasts, was loaded on each device and cells were seeded in each well by centrifugation. 2 millitissue devices were used per fibroblasts type. The organoids were imaged and fixed 3 days post-production with 4%PFA for 15 min at RT. Only tissues uniformly anchored to the tips of the cantilevers were included in further analysis.

### Immunostaining

A solution containing 2% bovine serum albumin (BSA), 2%FCS, 0.1% triton in PBS was used for permeabilization, blocking and dilution. Primary antibodies used in this study are: KRT14 (MA5-11599, ThermoFisher, 1:100), TBX20 (MAB8124, Novus Biologicals, 1:200), FOXA2 (ab108422, Abcam, 1:300), HHEX (MAB83771, R&D System, 1:100), FOXD1 (TA322737, OriGene, 1:50) PAX8 (NBP2-29903, Novus Biological, 1:100), TNT (RC-C2, DSHB 1:200). Cells were stained overnight at 4 °C, washed in PBS and incubated 1h with 1:500 secondary antibodies (Alexa Donkey anti Goat 568 - A11057, Goat anti Mouse 568 - A11031, Goat anti Rabbit 555 -A27017; ThermoFisher). Nuclei were counterstained with 0.1µg/ml DAPI (D1306, ThermoFisher).

For the immunostaining of Cardiac Microtissues, blocking and permeabilization were achieved with 0.1%Triton, 1% BSA in PBS (PBS-T-BSA) for 8 hours. The same solution was used to dilute primary antibodies: TNT (RC-C2, DSHB 1:200) or MF20 (DSHB 1:100). Staining was performed overnight under gentle agitation at 4 °C. After 3 x 5 min washes in PBS-T-BSA, microtissues were stained with secondary antibody and DAPI for 1h at room temperature. After staining, the PDMS devices were pulled out of the 35mm dish used as support and flipped on glass coverslip for confocal imaging.

### Imaging

Immunofluorescence images were acquired using either the either the upright fluorescent microscope Axio Imager.Z2 (Zeiss) or the SP8 confocal microscope equipped with a White Light Laser (Leica). For the cardiac microtissues, 3-4 tiles and 20-34 z stacks were imaged per sample. Tiles were combined using the LeicaX confocal software. Z-Stack projections and analysis were performed using Fiji version 1.0 and Imaris 8.4.1.

### Cell transfection and CRISPR knock-down

Cardiac fibroblasts from Rosa^Cas9-EGFP^ mice were transfected with guide RNAs using Lipofectamine MessengerMAX (ThermoFisher) according to manufacturer’s instructions. Briefly, 3 days post-isolation, CD45-CD31-CD90+ cells were FACS sorted and re-plated at about 10,000cells/cm^2^. After 6 days, when reaching 80-90% confluency, cells were incubated for 5’ with the RNA(1:50) -lipid(1:33) complex in Opti-MEM for 10’ at room temperature. Media was changed after 48h and cells were collected for RNA isolation at 72h. Two guide RNAs, designed and synthetized in-house (JAX Genetic Engineering Technologies facility), were used for each of the target genes (**Supplementary Table 7**). CleanCap® mCherry mRNA (TriLink Biotechnologies), and guide RNAs for GFP and scrambled guides were used as a controls. Guide RNA for the GFP gene were same as in [37].

### Cell transplantation in the kidney capsule

Adult fibroblasts from heart, tail and kidney were isolated from 10 weeks old Rosa ^mT/mG^ male mice as described above. After ten days cells were collected, counted and 4-5×10^5 cells were transferred in individual 1.5ml Eppendorf tubes (one per each kidney transplant), resuspended in 15-20ul of saline solution and kept on ice, until ready for the surgery. The remaining cells were re-plated (5×10^4/well of a mw6 plate) for the cultured cells controls. Syngeneic 10-11weeks old C57bl6/j mice were used for the surgeries. Mice were anesthetized with Tribromoethanol 400 mg/kg injected intraperitoneally. In the meanwhile, the cell suspension was spotted on a petri dish and fragments of sterile absorbable gelatin foam (Surgifoam, Ethicon), about 1mm long, were immersed in the drop.

Fur was removed from the left flank of the animal and eye ointment was applied. The mouse was placed in right lateral recumbency and a drape positioned over the surgical site. A 6-9 mm skin incision was made parallel and ventral to the spine and midway between the last rib and the iliac crest. A similar incision was made in the underlying abdominal wall. The kidney was externalized by placing forceps under the caudal pole and gently lifting through the incision and kept moist with warm sterile saline. A small incision was made in the capsule over the caudal-lateral aspect of the kidney, and a shallow subcapsular pocket was made with a blunt probe advanced toward the cranial pole of the kidney. The foam previously soaked in the cell suspension was placed in the far end of the subcapsular pocket. If needed, additional foam was used to close the incision site. Absorbable undyed 6.0 Vicryl sutures (Ethicon) were used to close the abdominal wall, 6.0 Vicryl black sutures (Ethicon) for the skin. Bupivacaine 0.1% was applied topically on the injection site and Slow-Release buprenorphine 0.05mg/kg was injected subcutaneously.

## Data availability

All transcriptome data that support the findings of this study are available in Gene Expression Omnibus (GEO) with the identifiers GSE98783 for microarray and SRR5590304 for single cell RNA sequencing. All other bulk RNAseq datasets generated for the CRISPR-experiment, kidney capsule transplant experiment and human cardiac biopsies comparison are available in GEO with the identifier GSE175765.

## Acknowledgements

We gratefully acknowledge the contribution of Luis E. Lima, Heidi Munger, Dr. Philipp Heinrich, and the Genome technology, Single Cell, Light Microscopy, Flow cytometry Services at The Jackson Laboratory for expert assistance with the work described in this publication.

This work is supported by grants from the Australian Research Council (ARC) and the National Health and Medical Research Council (NHMRC) to NAR, MWdC, MR, and NHMRC/Heart Foundation Fellowship to MR; and by the JAX Director’s Innovation Fund, and the NIH/NIGMS (2 P20 GM104318), the NIH/NIA (5 U01 AG022308-17), and grant from the Leducq Foundation for Cardiovascular Research to NR. The Australian Regenerative Medicine Institute is supported by grants from the State Government of Victoria and the Australian Government. The Jackson Laboratory scientific services are supported by the NIH/NCI (5 P30 CA034196).

## Author contributions

Conceptualization-MBF, EF, MR, HTN, NAR; Investigation-EF, MM, RC, SD, MWC; Data Curation-MBF, EF, MR, HTN, MWC; Formal analysis-MBF, EF, MR, HTN, MWC; Methodology-MBF, EF; Funding acquisition-MBF, EF, MR, HTN, NAR; Supervision-SB, TH; Project administration-MBF, EF; Manuscript writing MBF, EF, MR, HTN, NAR.

## Materials and correspondence

All material requests and correspondence should be directed to the corresponding author.

**Supplementary Fig. 1. Isolation of fibroblasts from different organs**. **a** Representative FACS plot showing the gating strategy used to isolate CD45-CD31-CD90+ fibroblasts from different organs. **b** Relative expression of markers *Vim, Thy1, Col1a2* in fibroblasts from gonads, heart, kidney, liver, lung, and skin, compared to tail. **c** Raw Ct values for *Vim, Thy1, Col1a2* and *Hprt1* qPCR analysis. Data are mean ± SEM on 3 biological replicates.

**Supplementary Fig. 2. Generic Molecular Signature of Organ Fibroblasts. a** High signal intensity gene of all fibroblast samples without relative quantification were entered in IPA and molecular pathways were generated, revealing common processes among all fibroblasts. Most genes were involved in cell death and survival, cell signaling and molecular transport and trafficking. **b** High intensity genes organized by cell compartment. The microarray analysis was performed on 3 biological replicates per each fibroblasts type.

**Supplementary Fig. 3. Pairwise comparison between the Hox code of organ fibroblasts.** Related to Fig.1. Heat map plot showing the pairwise square Euclidean distances based on raw signal intensity of organ fibroblasts (n=3). Smaller distance means the two organ fibroblasts are transcriptionally more similar.

**Supplementary Fig. 4. Embryological Molecular Signature of Organ Fibroblasts.** Related to Fig.2. Additional validations of the expression of some organ-enriched, developmental related genes using qPCR, on adult and embryonic derived fibroblasts. Data are mean ± SEM on 3 biological replicates.

**Supplementary Fig. 5. Molecular Signature of Skin Fibroblasts. a** Heatmap highlighting the genes differentially expressed over 10-fold solely in skin fibroblasts. **b-f**, IPA analysis on genes highly expressed in skin fibroblasts (shown in **a**). **b** top 5 canonical pathways with p-value and estimated percentage of overlap**. c** Top diseases and biological functions. **d** Top networks and associated functions. **e** representation of a pathway related to skin embryonic development (Morphogenesis of epithelial tissue) with related skin-fibroblasts enriched genes. **f** graphic representation of the top network (highlighted with a red arrow in **d**) overlaid with the expression of genes enriched in our dataset (circled in pink) associated with skin disease and function. Red arrows point to all skin/derma related terms.

**Supplementary Fig. 6. Molecular Signature of Lung Fibroblasts. a** Heatmap highlighting the genes differentially expressed over 10-fold solely in lung fibroblasts. **b-f**, IPA analysis on genes highly expressed in lung fibroblasts (shown in **a**). **b** top 5 canonical pathways with p-value and estimated percentage of overlap**. c** Top diseases and biological functions. **d** Top networks and associated functions. **e** representation of a development associated pathway (Respiratory system development) with related lung-fibroblasts enriched genes. **f** graphic representation of the top network (highlighted with a red arrow in **d**) overlaid with the expression of genes enriched in our dataset (circled in pink), associated with lung development and function.

**Supplementary Fig. 7. Molecular Signature of Liver Fibroblasts. a** Heatmap highlighting the genes differentially expressed over 10-fold solely in liver fibroblasts. **b-f**, IPA analysis on genes highly expressed in liver fibroblasts (shown in **a**). **b** top 5 canonical pathways with p-value and estimated percentage of overlap**. c** Top diseases and biological functions. **d** Top networks and associated functions. **e** representation of a development associated pathway (Development of liver) with related liver-fibroblasts enriched genes. **f** graphic representation of the top network (highlighted with a red arrow in **d**) overlaid with the expression of genes enriched in our dataset (circled in pink), associated with liver abnormal development and function.

**Supplementary Fig. 8. Molecular Signature of Kidney Fibroblasts. a** Heatmap highlighting the genes differentially expressed over 10-fold solely in kidney fibroblasts. **b-f**, IPA analysis on genes highly expressed in kidney fibroblasts (shown in **a**). **b** top 5 canonical pathways with p-value and estimated percentage of overlap**. c** Top diseases and biological functions. **d** Top networks and associated functions. **e** top toxicology lists, including different list of genes associated with kidney injury. **f** representation of development associated pathways (Development of metanephric mesenchyme, metanephros, formation of kidneys) with related kidney-fibroblasts enriched genes. **g** graphic representation of the top network (highlighted with a red arrow in **d**) overlaid with the expression of genes enriched in our dataset (circled in pink), associated with kidney disease.

**Supplementary Fig. 9. Molecular Signature of Gonad Fibroblasts. a** Heatmap highlighting the genes differentially expressed over 10-fold solely in gonad (testis) fibroblasts. **b-f**, IPA analysis on genes highly expressed in gonad fibroblasts (shown in **a**). **b** top 5 canonical pathways with p-value and estimated percentage of overlap**. c** Top diseases and biological functions. **d** Top networks and associated functions. **e** representation of development associated pathways (morphology of genital organs, sex determination, size of genital organs), with related gonad-fibroblasts enriched genes. **f** graphic representation of the top network (highlighted with a red arrow in **d**) overlaid with the expression of genes enriched in our dataset (circled in pink), associated with reproductive system development and disfunction.

**Supplementary Fig. 10. Molecular Signature of Cardiac Fibroblasts. a** Heatmap highlighting the genes differentially expressed over 10-fold solely in cardiac fibroblasts. **b-f**, IPA analysis on genes highly expressed in cardiac fibroblasts (shown in **a**). **b** top 5 canonical pathways with p-value and estimated percentage of overlap**. c** Top diseases and biological functions. **d** Top networks and associated functions. **e** representation of development associated pathways (Development of pericardium, hyper-trabeculation, innervation, hypoplastic heart syndrome) with related heart-fibroblasts enriched genes. **f** graphic representation of the second top network (highlighted with a red arrow in **d**) overlaid with the expression of genes enriched in our dataset (circled in pink), associated with cardiac diseases and disorders. **g** Heatmap showing the expression of cardiac fibroblasts-enriched genes in human left ventricular biopsies from healthy (N=5) and chronic ischemic heart failure patients (N=5).

**Supplementary Fig. 11. Analysis of the organ specific fibroblasts heterogeneity at single cell level.** Related to Fig.4. Data derived from the re-analysis of Mouse Cell Atlas stromal cell dataset. **a** Heatmap showing the top differentially expressed genes in each cell of the sub-cluster Lung A-B-C. The genes were identified by pairwise differential expression of Lung C versus Lung A, LungC versus Lung B and Lung B versus Lung A. **b** Heatmap showing the average expression per sub-population of key-defining gene and the lung-development related genes (*Foxf1*). **c** Heatmap showing the top differentially expressed genes in kidneyA-B. The genes were identified by pairwise differential expression. **d** Heatmap showing the average expression per sub-population of key defining genes and the kidney-development related genes (*Pax8, Wnt7b, Bmp7*).

**Supplementary Fig. 12. Analysis of the organ specific fibroblasts heterogeneity at single cell level.** Related to Fig.4. Data derived from cultured fibroblasts scRNAseq data generated in this manuscript. **a.** Heatmap showing top differentially expressed genes between Heart 1 and 2, **b** Heatmap showing the average expression per sub-population of key defining genes and the heart-development related genes (*Itga4, Col2a1, Tbx20*). **e.** Heatmap showing the expression of genes identified by pairwise differential expression analysis of freshly isolated kidney fibroblasts (same as in **Supplementary Fig. 8c**), **f** Heatmap showing top differentially expressed genes between Kidney 1 and 2, **g** Heatmap showing the average expression per sub-population of key defining genes and the kidney-development related genes (*Pax8, Wnt7b, Bmp7*).

**Supplementary Fig. 13. Analysis of the fibroblast’s specific response to transplant under the kidney capsule.** Related to Fig.7. **a.** Table showing the genes associated to the top KEGG pathways in Fig 7m, **b** Canonical pathways identified through Ingenuity Pathway Analysis of the differentially expressed genes, ordered by significance (-LOG of the B-H p-value) and colored by the activation z-score predicted for the three comparisons HFs/HFc, TFs/TFc, KFs/KFc.

**Supplementary Table 1. Microarray data: highly expressed genes common to all organ-specific fibroblast populations, classified based on cellular process or cellular localization.**

**Supplementary Table 2. Microarray data: average raw expression and standard errors of Hox code genes across all fibroblast samples from the microarray analysis (n=3).**

**Supplementary Table 3. Microarray data: expression of genes that were enriched by 10-fold change or more in single organ fibroblasts compared to tail fibroblasts (n=3).**

**Supplementary Table 4. Expression of cardiac fibroblasts enriched genes in human left ventricular biopsies from healthy and chronic ischemic heart failure patients.**

**Supplementary Table 5. Analysis of the stromal cell aggregate from the Mouse Cell Atlas: markers genes per each population, and markers identified by pairwise comparison of the 2 kidney and 3 lung populations.**

**Supplementary Table 6. Analysis of in-house scRNAseq data of merged cultured fibroblasts from different organs: markers genes per each population, and markers identified by pairwise comparison of the 2 kidney and 2 cardiac populations.**

**Supplementary table 7: CRISPR-Cas9 experiments: sequence of the guide RNAs; differential expressed genes between Tbx20 and Gata4 KD and the correspondent controls. KEGG pathways analysis.**

**Supplementary Table 8. Sequence of all the qPCR primers used in the study.**

**Supplementary video 1. Co-culture of adult cardiac fibroblasts with neonatal ventricular cardiomyocytes. 20x magnification.**

**Supplementary video 2. Co-culture of adult kidney fibroblasts with neonatal ventricular cardiomyocytes. 20x magnification.**

